# A Geometrical Optics Model of Individual Ommatidia in Compound Eyes and a Hypothesis for Monocular Stereopsis

**DOI:** 10.1101/2025.06.19.660517

**Authors:** Yong Ding, Dalu Ding, Dawei Ding, Rongzhi Jiang

## Abstract

This study proposes a novel geometrical optics-based visual model suggesting that individual ommatidia in insect compound eyes may possess intrinsic three-dimensional imaging capabilities and enable monocular stereoscopic perception. Building upon the authors’ previous hypothesis that certain vertebrate retinal cell nuclei act as microlenses, the optical model is extended in this work to the arthropod compound eye system. By conceptualizing the cornea, crystalline cones, and layered rhabdomeric nuclei as components of a “multi-angle miniature real-image telescope,” we construct a three-tiered imaging system composed of objectives, relay conjugate lenses, and microlenses. The model predicts that an ommatidium can actively perceive spatial depth and object distance by adjusting the focal length of crystalline cones and the distance between focused planes, thus achieving stereoscopic imaging with a single eye. Simulations based on representative species such as *Drosophila melanogaster, Viggiani* parasitic wasps, and fireflies (*Lampyris*) support the feasibility of this model in distance perception across scales ranging from microns to meters. The model breaks away from the “one ommatidium—one image point” paradigm and offers a new theoretical architecture for understanding insect stereopsis. It also presents a promising framework for bioinspired 3D imaging systems and optical designs with multi-focal capabilities.

## 1. Introduction

The mechanisms underlying biological vision remain a central theme in optics and neuroscience. Conventional wisdom distinguishes the optical architectures of vertebrate eyes and arthropod compound eyes: the former relies on a unitary lens system (cornea-lens complex) for image formation, while the latter operates via multiple spatially distributed ommatidia. These paradigms, however, typically overlook the potential active optical contributions of internal subcellular structures—particularly cell nuclei—within the retina or individual ommatidium. Whether photoreceptor or interneuron nuclei could participate directly in 3D image formation remains an open, underexplored question. In late 2024, a groundbreaking hypothesis was presented by the authors [1,2], proposing that neural cell nuclei within the vertebrate retina—including those of ganglion cells, bipolar cells, and rod/cone photoreceptors—function as microlenses contributing to real image formation. On this theoretical foundation, a new optical model for single-eye imaging was built, centered on the concept of the retina as a high-resolution microlens array with angular diversity.

### 1.1 Hypothesis: “Array of High-Magnification Miniature Telescopes” in Single-Eye Vision

Contrary to the traditional view where the eye uses a dominant lens (crystalline lens in humans), this model posits an eye as a multi-microlens system. Each microlens telescope operates with its own azimuth and elevation angles. The imaging sequence includes:

1. **Objective lens system:** The cornea and lens collaborate to form a primary aerial image.
2. **Conjugate relay system:** Neural cell nuclei (ganglion and bipolar cells) form a conjugate mirror array, relaying and refocusing the image.
3. **Terminal depth detection system:** Photoreceptor cell nuclei act as microlenses, real-imaging onto the outer segments of their respective photoreceptors.

Brain analysis circuitry then determines spatial depth based on image clarity and photic signal intensity, adjusted through changes in lens focal length and lens-conjugate-microlens spacing.

### 1.2 Hypothesis: “Four-Angle Miniature Real-Image Telescope” within an Ommatidium

Extending this concept into insect compound eyes, the study proposes that each ommatidium—with a cornea, crystalline cones, and rhabdomere nuclei—functions similarly to a four-angle stereoscopic imaging telescope. Under the assumption that cone and rhabdom nuclei act as microlenses, each unit includes:

1. **Four-angle objective system:** One cornea + four crystalline cones imaging at different angular positions (eight positional vectors total).
2. **Conjugate relay system:** Six R-cell nuclei (rhabdom nuclei), layered above and below, act as focusing intermediates.
3. **Terminal depth perception system:** Two to three rhabdom nuclei at distinct axial depths act as microlenses creating sharp final images on different rhabdom bundles.
4. **Miniature depth sensors:** Rhodopsin-containing segments in rhabdoms convert light into electrical signals, each tuned to a specific depth focus. By modulating cone focal length or changing focal plane separation, these systems may detect object distances. Modeling and optic simulation based on anatomical parameters of three diverse species—*Drosophila, Viggiani* wasps, and fireflies (*Lampyris*) —support the viability of this stereoscopic model at micro-to meter scales.

## 2. Geometrical Optics Model of Individual Ommatidia, Hypothesis of Stereoscopic Perception Mechanism, Derivation of Distance Estimation Equations, and Numerical Simulations

### 2.1 Hypothetical 3D Imaging Model of a Single Ommatidium: “Four-Angle Miniature Real-Image Telescope”

This study hypothesizes that the rhabdomeric nuclei within an ommatidium possess microlens properties, hereafter referred to as *rhabdom nucleus microlenses*. Based on this foundational assumption, and referencing anatomical data from *Viggiani* parasitic wasps, *Drosophila*, and fireflies (*Lampyris*) [3–8], we present schematic diagrams (Figure 1 A– C) illustrating the proposed 3D optical imaging model of a single ommatidium. To simplify the presentation of the spatial geometry, we illustrate the system in two orthogonal optical meridional planes (Figures 1A and 1B, corresponding to elevation angles ϕ = 0° and ϕ = 90° respectively, within a spherical coordinate system). Although seven or eight rhabdom nuclei are spatially distributed in 3D, they are projected onto these two cross-sectional views for clarity.

**Figure 1.**
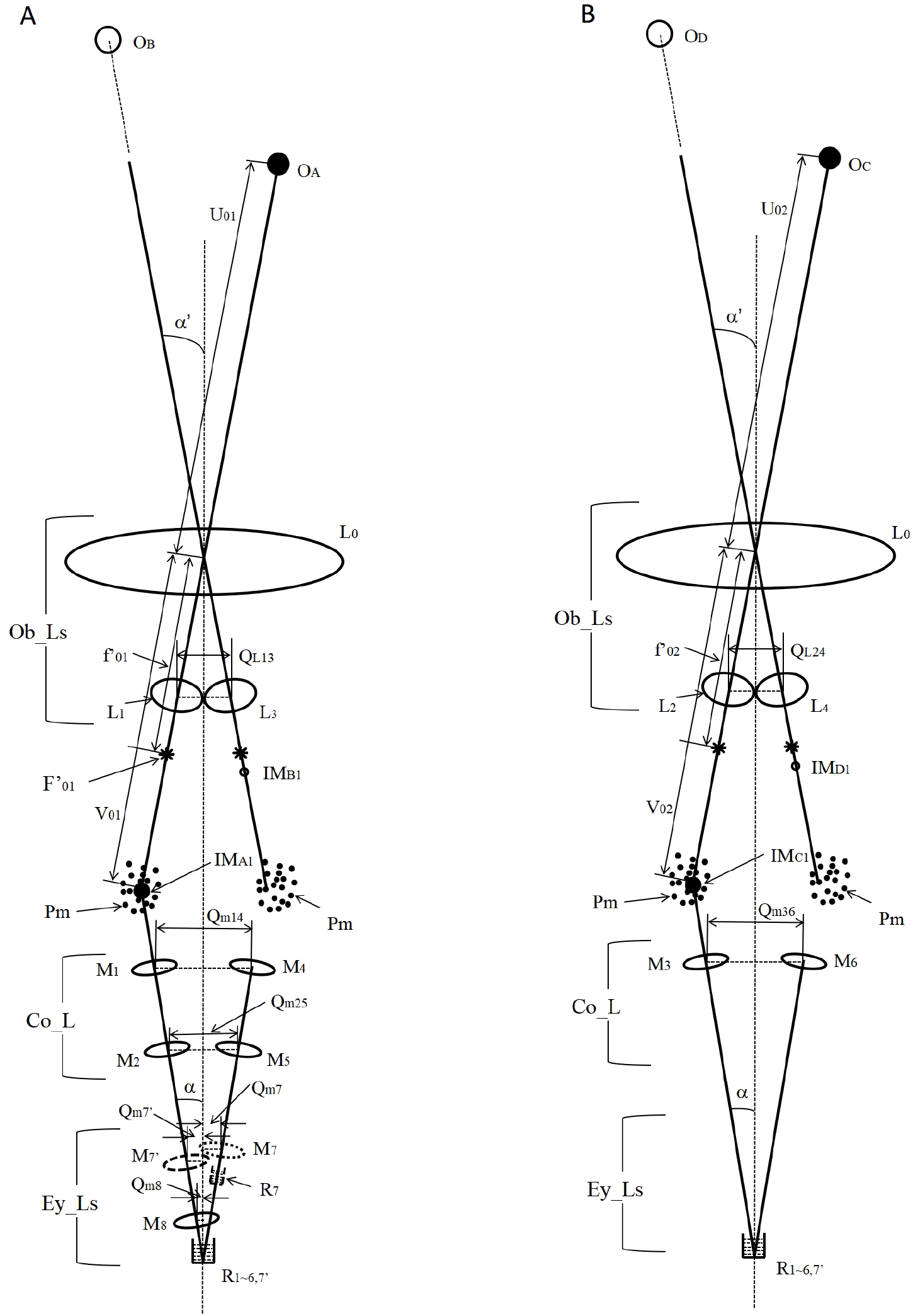
Schematic representation of the 3D optical imaging model within a single ommatidium. This figure illustrates the proposed three-tier real-image optical model of a single ommatidium conceptualized as a “four-angle miniature real-image telescope.” Components are labeled as follows: •**Ob_Ls**: Objectives (formed by cornea L0 + one of four crystalline cones L1–L4) •**Co_L**\*\*: Conjugate (relay) lens system formed by six rhabdomeric nuclei (M1–M6) •**Ey_Ls**: Microlens system comprising M7, M7′, M8 at differing depth layers •**Pm**: Pigment cells •**L0–L4**: Cornea (L0) and four crystalline cones (L1–L4), each forming an objective •**Objectives**: Each objective lens is composed of L0 (cornea) and one of L1–L4 •**M1–M6**: Rhabdom nuclei forming a conjugate mirror system (relay optics) •**M7, M7′, M8**: Base-level rhabdom nuclei functioning as terminal microlenses •**R7, R7′, R8**: Corresponding rhabdom bundles functioning as mini-depth sensors •**QL13, QL24**: Distance between centers of L1–L3 and L2–L4 (i.e., crystalline cones) •**Qm14, Qm36, Qm25**: Center-to-center distances between rhabdom nuclei — M1 & M4, M3 & M6, M2 & M5, respectively •**Qm7, Qm7′, Qm8**: Vertical distances from M7, M7′, and M8 to the ommatidial central axis •**OA, OB, OC, OD**: Spatial object points A, B, C, and D •**IMA1–IMD1**: First image points corresponding to object points OA–OD •**IMA2, IMA3**: Second and third image points; IMA2 is the conjugate image of IMA1, and IMA3 is the image of IMA2 formed by the microlens •**\***: Denotes object-side or image-side focal points of the microlenses •**α**: Indicates either the angle between the optical axis of a microlens (e.g., M7′) and the central axis of the ommatidium, or the inner/outer semi-field angle of the ommatidium •**f′**_**01**_, **f′**_**02**_: Image-side focal lengths of the objective lenses formed by L0–L1 and L0–L2 respectively •**F′**_**01**_: Image-side focal point position for the L0–L1 objective lens •**F′\***_**01**_ **(or f′\***_**01**_): Conjugate focal point (or conjugate focal length) corresponding to F′_01_ •**U**_**01**_, **U**_**02**_: Object distances for object points seen through the L0–L1 and L0–L2 objective lenses •**V**_**01**_, **V**_**02**_: Image distances corresponding to U_01_ and U_02_ •**Um1, Um4, Um2, Um5**: Object distances or conjugate object distances for relay nuclei M1, M4, M2, M5 •**Vm1, Vm4, Vm2, Vm5**: Image distances corresponding to those nuclei •**Df (Δfocal)**: Distance between the conjugate focal plane of the image-side of the objective (after passing through the relay) and the object-side focal plane of the terminal microlens (M7, M7′, M8)

#### Key components of the optical system are

1. **Objective Lenses (n = 4):**Each formed by the cornea and one of four crystalline cones (L1–L4), each with distinct angular orientations (azimuth and elevation). These create primary images (first image points) for respective angular segments. First image points (IMA1, IMB1, etc.) are formed from spatial objects (OA, OB, etc.) as seen in Figures 1A and 1B.
2. **One Conjugate Mirror:**Composed of six rhabdomeric nuclei (M1, M4, M3, M6 in the upper layer, and M2, M5 in the lower layer), acting as relay lenses. These nuclei re-image the first focal points to create conjugate second images (e.g., IMA2 from IMA1), as shown in Figure 1C.
3. **Microlens Eyepices at Ommatidial Base:**Three rhabdomeric nuclei (M7, M7’, M8) embedded near the ommatidial base serve as microlenses. They project real third images of the conjugate image onto the distal rhabdom bundles (e.g., R7, R7’, R8). Crucially, for real image formation, the conjugate image lies outside their focal range, unlike traditional telescopes.
4. **Microscale Depth Detectors:** The rhabdoms R7, R7’, and R8 function as *miniature depth detectors*, converting focused light at different depths into phototransduction signals (i.e., electrical impulses). Each microlens-rhabdom pair has a distinct operable range of object distances.
5. **Light Scatting Pigments :** Apart from blocking stray light, pigments help redirect scattered light from each image level toward subsequent imaging elements, enhancing the efficiency of the optical chain. An analogous function has been proposed for yellow pigments in the human fovea [1,2].

Under this theoretical construct, a single ommatidium can be treated as a **four-angle miniature stereo-imaging telescope**, capable of perceiving a target’s angular position and spatial depth within the visual field.

### 2.2 Hypothesis: Perception Process of Depth and Stereopsis by a Single Ommatidium

To model the hypothesized stereoscopic depth-sensing mechanism, we describe the imaging process per Figure 1 (A–C), under the following simplifying assumptions:

1. All four objective lenses (formed by L0-L1 to L4) share the same focal length. Similarly, all rhabdom nucleus microlenses possess identical focal parameters.
2. The optical conjugate plane of the relay lens (mirror) always coincides with or lies within the conjugating depth of the first image plane from the objectives. Hence, a clear second (relay) image is always achieved.
3. For each rhabdom nucleus microlens, the effective image distance at which rhabdoms detect steep increases in signal (i.e., peak phototransduction) is fixed and thus corresponds to a unique object distance.
4. We define **focal plane separation (Df)** as the distance between the conjugate focal plane of the objective-cone system (after the relay) and the object-side focal plane of the rhabdom nucleus microlens system.
5. The insect brain computes the object distance based on the value of **object focal length (f0i or f’0i)** or the **focal plane separation (Df)** at which the photocurrent increase occurred (possibly related to the muscular tension required to adjust the cone lens array [9,10]).

#### Two sensory mechanisms are proposed based on the direction of spatial scanning

1) **From Near to Medium-Distance Perception (Df = 0)**

- The system operates by scanning through focal length variation in the objective (i.e., adjusting cone length/refractive power) while keeping Df = 0.
- As a spatial object projects light into the ommatidium, real images sequentially pass through rhabdom layers. When the image depth coincides with a specific rhabdom bundle, a *sharp spike in photoreceptor current* is expected.
- The insect’s neural system then recognizes the object distance based on:
  ▪ The focal length setting at the sharp signal onset
  ▪ The specific rhabdom microlens involved in the spike
- Rhabdoms M8, M7’, M7 are considered successively responsible for near, mid, and far-range detections.

2) **From Medium-Distance to Infinity (Df > 0)**

- With the objective lens focal length fixed at its maximum, the system increases focal plane separation Df to iteratively probe further depths.
- The microlens M7 is primarily involved in this range.
- A photoreceptor at M7’s rhabdom will signal a sharp current increase when a conjugate image aligns with its focus across varying Df values.
- The brain again interprets distance from:
  ▪ The triggered Df value
  ▪ The identity of the activated rhabdom microlens

This modular design allows the detection of object distances across overlapping ranges, using rhabdom depth stratification and differential optical path lengths. The mechanism essentially parallels a variable-focus stereoscope, functioning within a single ommatidium.

### 2.3 Derivation of Object Distance Estimation Equations in a Single Ommatidium

In the following derivation, all symbol definitions (e.g., e.g., *Umi, Vmi, fmi* , representing the object distance, image distance, and focal length of microlens Mi, respectively) are consistent with the annotations provided in Figure 1. Note that object and image distances for both the objective and microlens systems follow standard geometric optics sign conventions [11,12], where object-side distances are treated as negative.

Based on the stereoscopic perception mechanism proposed in Section 2.2, and focusing on the imaging pathway involving microlens M7 (with a fixed image distance), we derive the forward imaging equation along a specific optical path: from the main objective lens (formed by cornea L0 and crystalline cone L1) to the rhabdom beneath microlens M7.

The derivation is grounded in the principles of sequential lens imaging and aims to calculate the object distance *U01*.

The imaging equation for microlens M7 is:

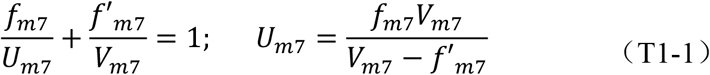

The distance between the possitions of object point and object side focal point of microlens M7 is:

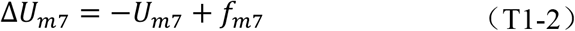

The distance between the object lens’s conjugate focal point *F*^’*^_01_ and the center of microlens M2 is:

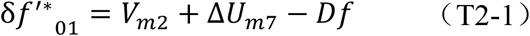

According to conjugate imaging principles, the distance between the object lens’ image side focal point *F*^’^_01_ and the center of microlens M1 is:

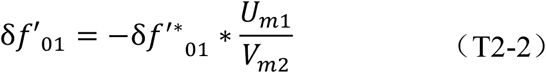

The distance between the object lens’ image side focal point and object point of microlens M1 is:

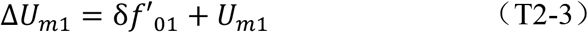

The distance between the the object lens’ image and foccal points (assuming the focal plane of the ojective lens L0+L1 is spherical):

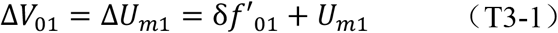

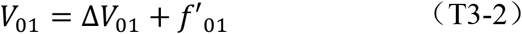

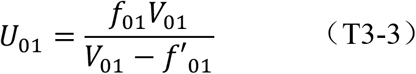

These equations demonstrate that when the following parameters are fixed:

- Focal lengths of the micro-eyepiece and microlenses for the conjugate mirror
- Object and image distances of the micro-eyepiece
- Conjugate distance of the conjugate mirror

Then, the object distance *U*01 of a spatial point is determined by the **focal length of the objective lens *f***_**01**_ **or *f*’** _**01**_ and the **focal plane spacing *Df***.

#### Supplemental Derivation for the Conjugate Lens Imaging Process (Figure 1C Reference)

We further illustrate how the conjugate lens system forms the second image (IMA2) from the first image (IMA1) through microlenses M1=M6. Using trigonometric relations and conjugate imaging formulas, we define:

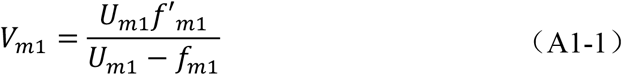

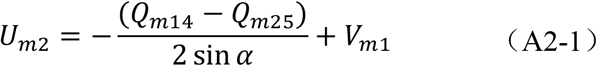

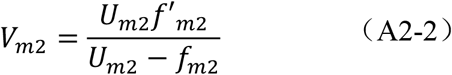

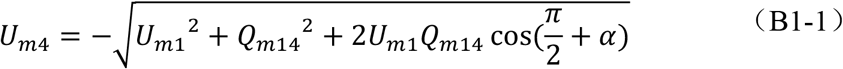

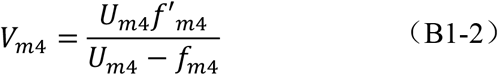

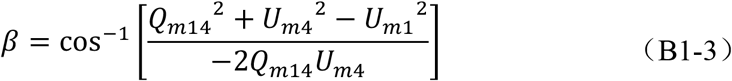

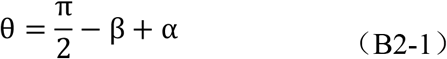

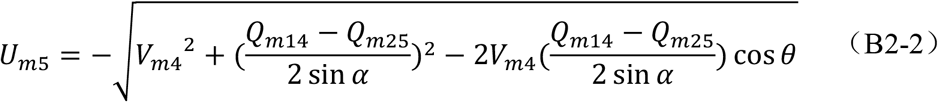

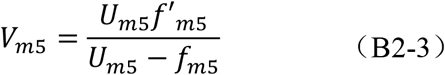

Equations A1-1 to A2-2: Determine the imaging process through microlenses M1 and M2 and their geometric relations.

Equations B1-1 to B2-3: Determine the imaging process through microlenses M4 and M5 and their geometric relations.

To satisfy the condition of conjugate imaging, the images formed by M2 and M5 must overlap or coincide. Thus, they must also satisfy:

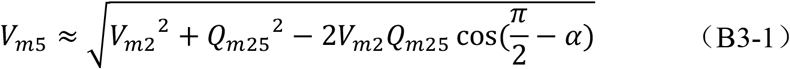

When the relative positions and focal lengths of microlenses within the conjugate system are fixed, the two sets of equations (A and B) show that the same spatial point can produce a consistent conjugate image through multiple microlenses, leading to **multiple valid optical paths**. These discrete but equivalent solutions allow the model to be robust to optical variation and noise.

### 2.3 Numerical Simulation of Depth Perception by Single Ommatidia

#### 2.4.1 *Viggiani* Parasitic Wasp Simulation

Using anatomical metrics derived from literature [3,4], Table 1 shows estimated optical parameters of the wasp’s ommatidium model, assuming an internal refractive index n = 1.33 and uniform focal length among rhabdom nuclei microlenses.

**Table 1.**
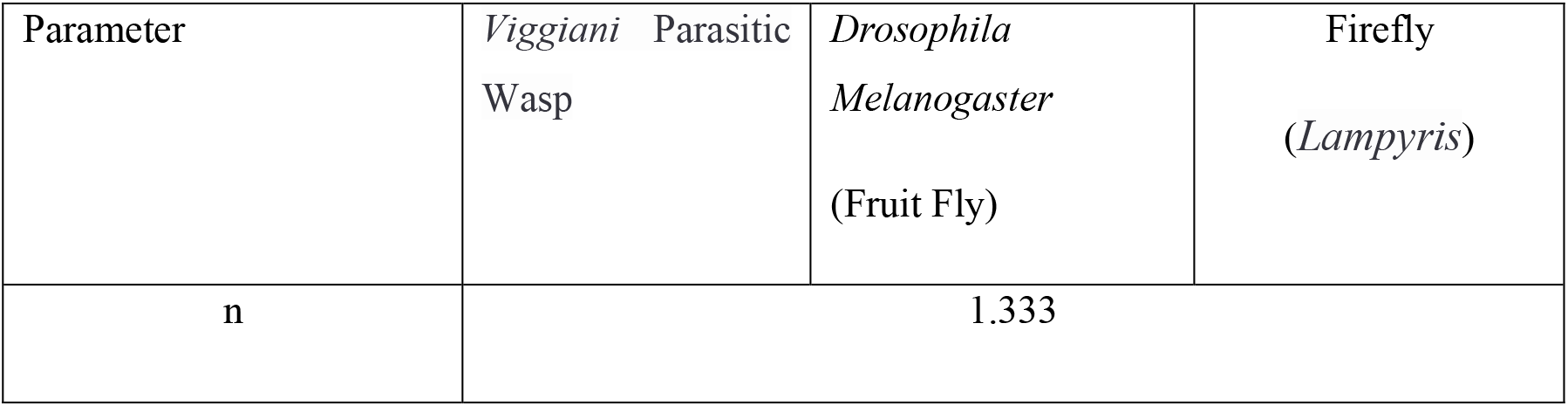
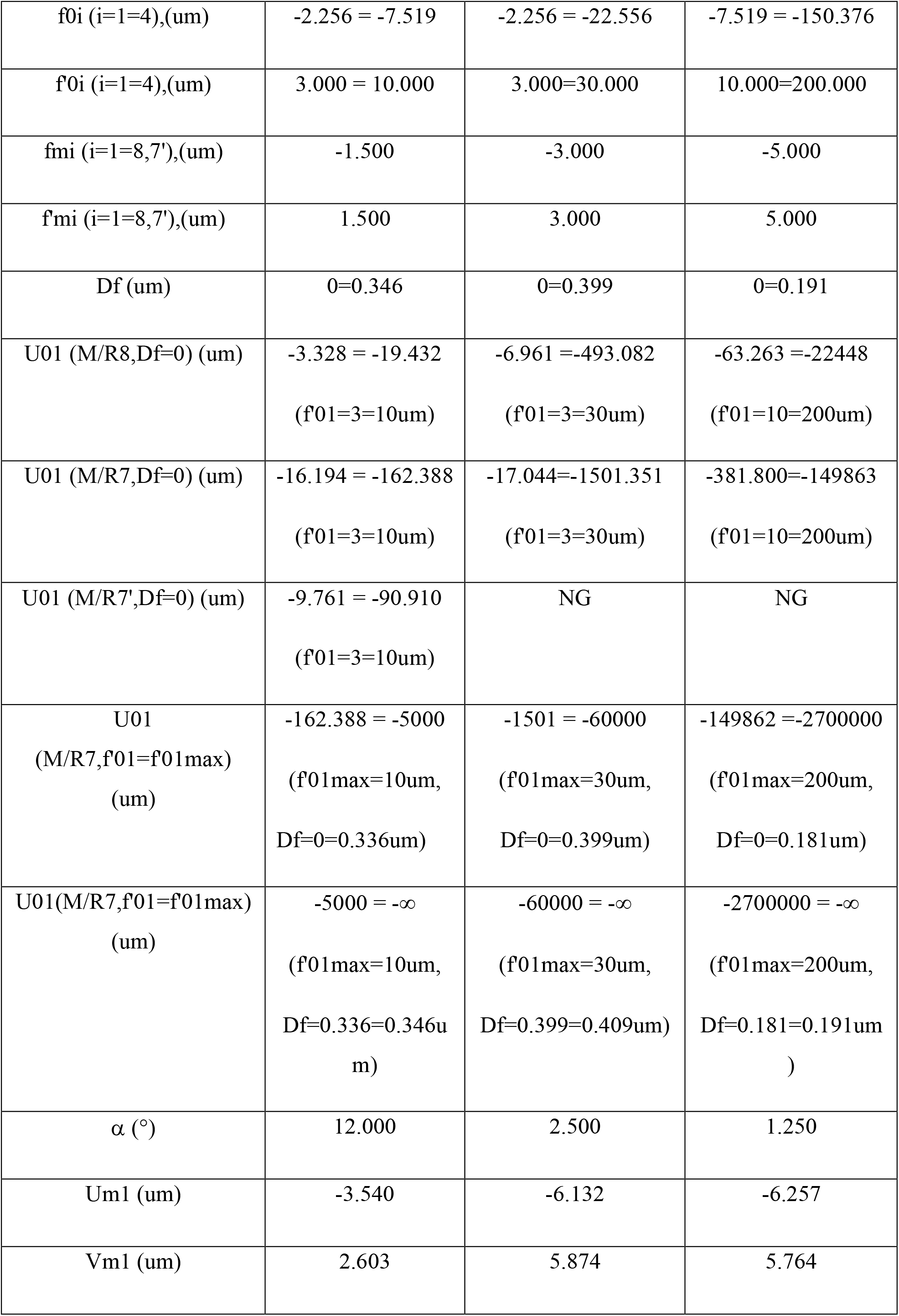
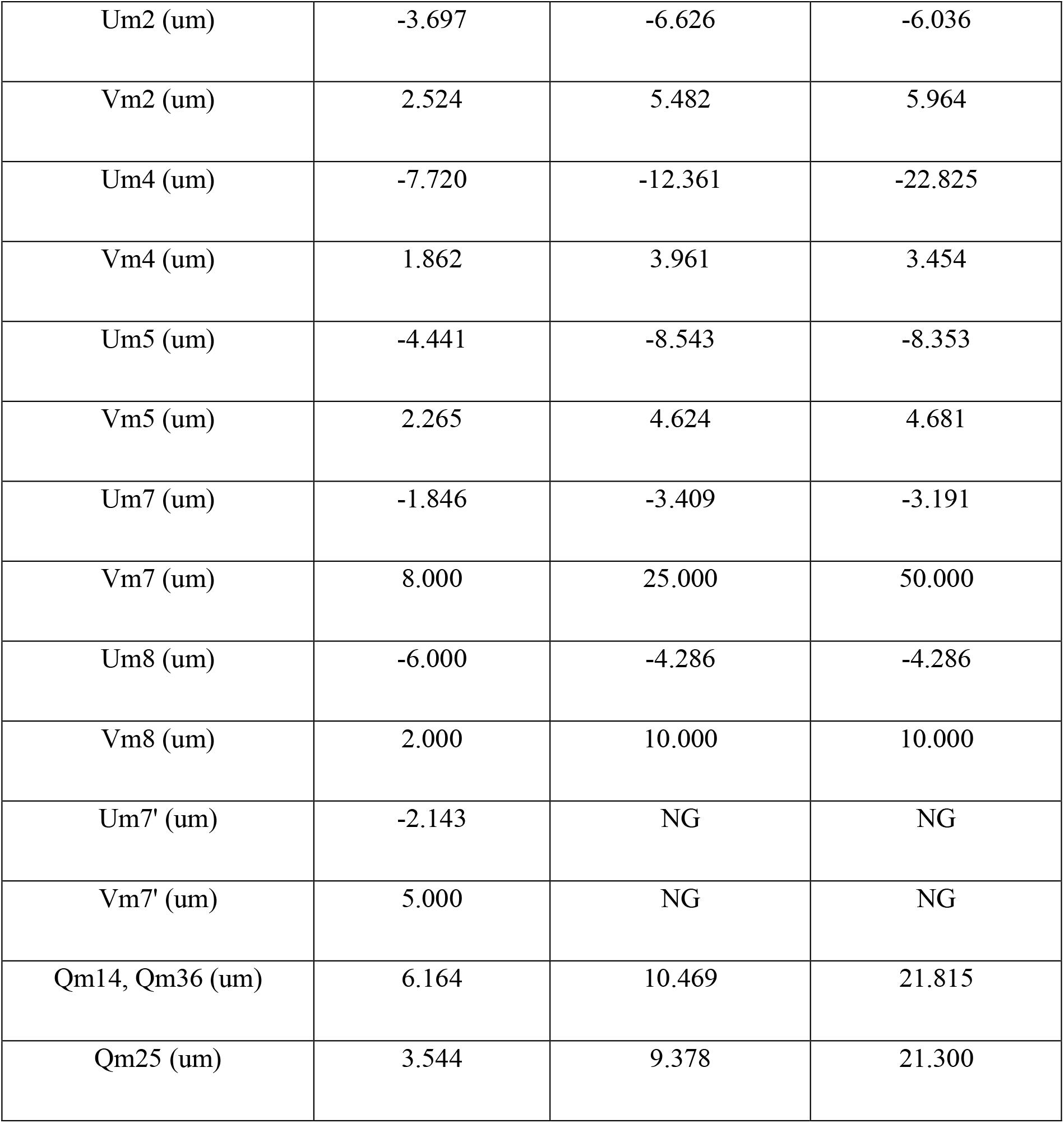
Estimated Geometric Optical Parameters of Microlenses in the 3D Imaging Model of a Single Ommatidium.

#### Key derived parameters (abbreviated)

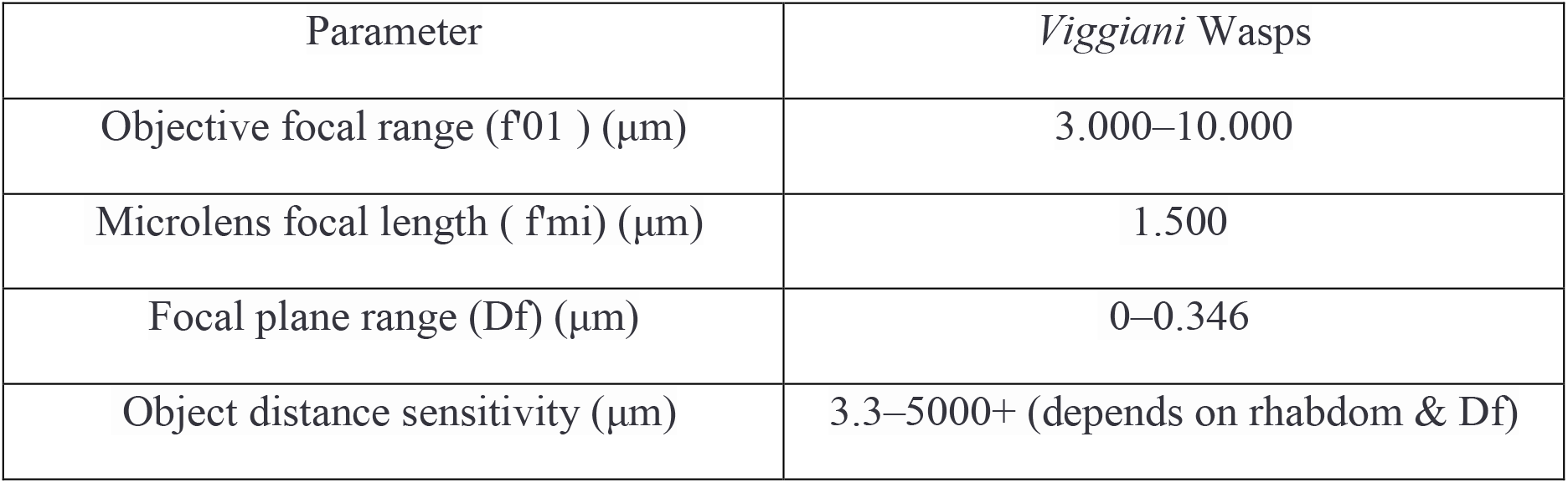

#### Results

##### Figure 2

When Df = 0, by scanning the objective focal length f’01 from 3–10μm, object distances that sharply focus on different rhabdom bundles are:

**Figure 2.**
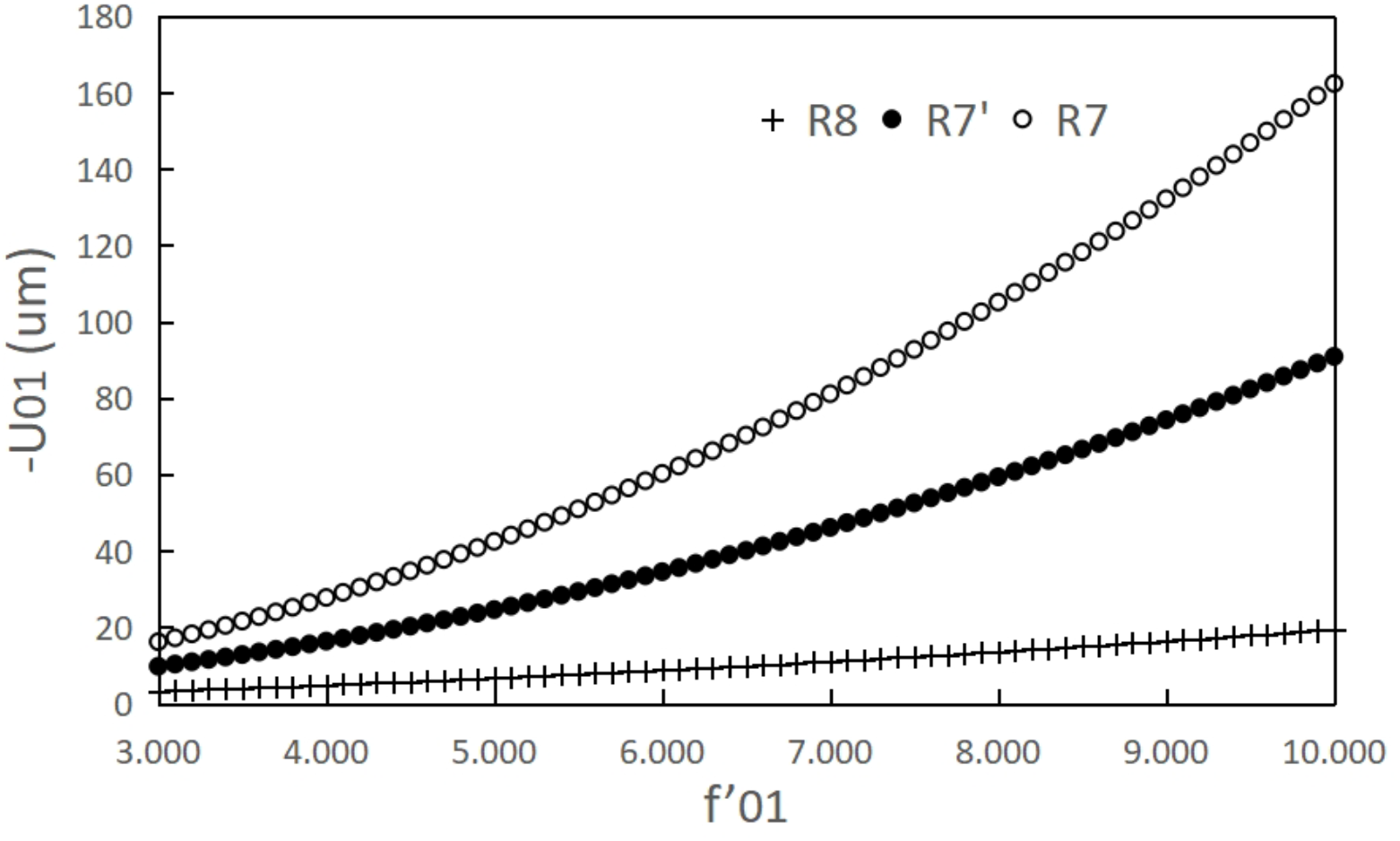
Simulation of object distance perception range in *Viggiani* parasitic wasps when Df = 0. The relationship between objective focal length f’0i= 3–10 μm and estimated object distance is plotted for rhabdom focal points R8, R7′, and R7. The corresponding perception ranges for each microlens are approximately: - R8: 3.3–19.4 μm - R7′: 9.8–90.9 μ - R7: 16.2–162.4 μm These results demonstrate that precise object depth detection is feasible using only objective focal length modulation.

‐ R8: 3.3–19.4μm
‐ R7′: 9.8–90.9μm
‐ R7: 16.2–162.4μm

##### Figure 3

Fixing f’01 = 10μm and varying Df from 0 to 0.336μm:

**Figure 3.**
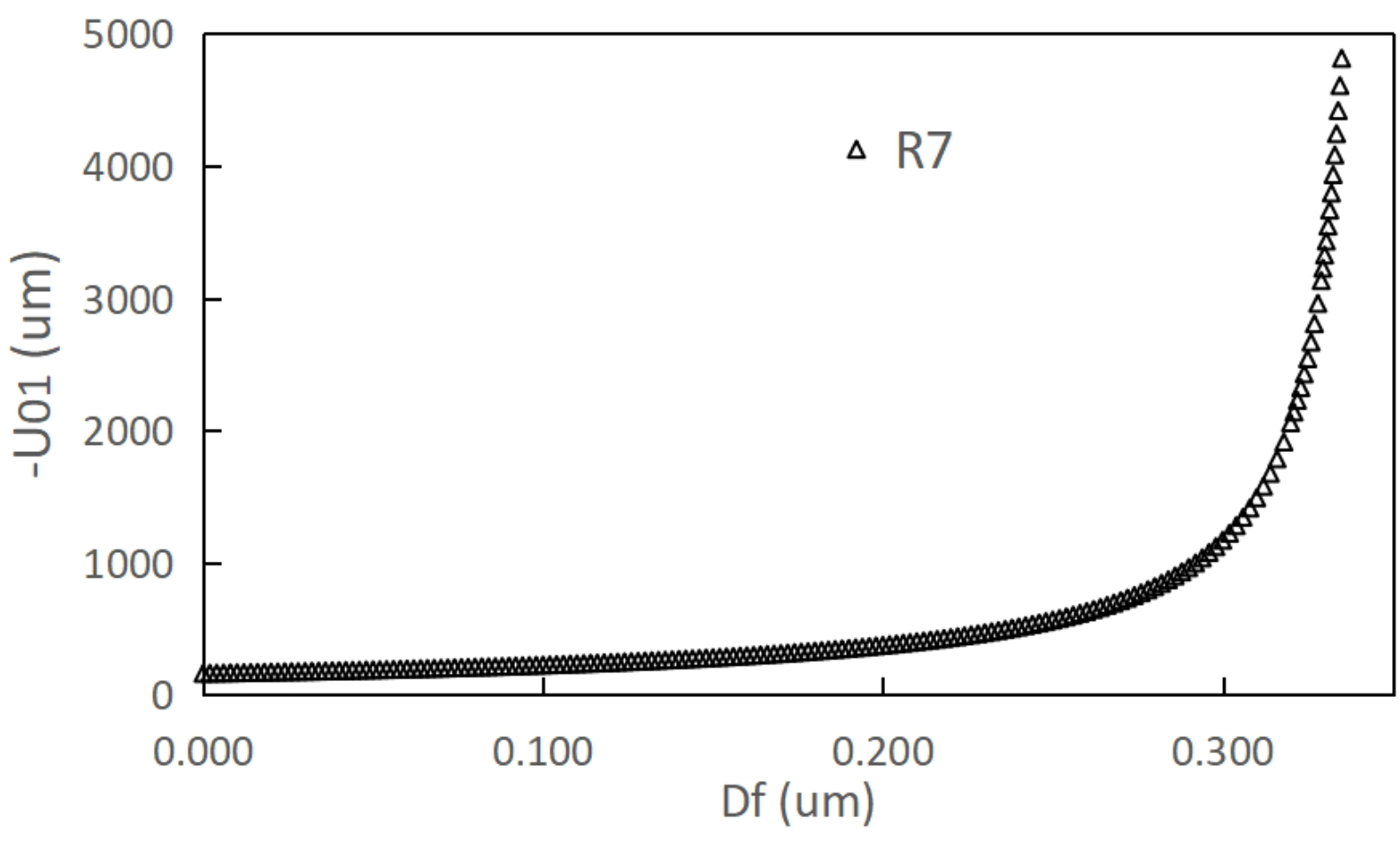
Object distance vs. focal plane spacing (Df) simulation in *Viggiani* wasps when objective focal length is fixed. With maximum objective focal length f’0imax= 10μm, increasing focal plane spacing Df = 0– 0.336μm enables the microlens M7 to detect object distances from 162 μm to over 5,000 μm. Beyond Df > 0.346μm, objects appear optically at infinity. The system demonstrates capability to resolve micro- to millimeter depth.

‐ Detected object distance increases from 162.4μm to 5000μm
‐ Df > 0.346μm corresponds to object distances approaching ∞

### Conclusion

With a focal plane modulation precision of 10 nm, *Viggiani* wasps can distinguish depth differences within <5 mm, but lose distance discrimination precision beyond that range — object presence is detected, but not exact range.

#### 2.4.2 *Drosophila* (Fruit Fly) Simulation

Based on EM studies in [5,6], simulations identified visual range expansions in flies.

Parameter Highlights:

‐ Objective f’0i: 3–30μm
‐ Df range: 0–0.399μm
‐ R7 detection range (Df = 0): 17–1500μm
‐ R7 with increasing Df: 1.5 mm – 60 mm
‐ Beyond Df > 0.409 μm: infinite range anticipation

Thus, *Drosophila* may discern distance as fine as millimeter-resolution up to 6 cm using a single ommatidium.

**Figure 4.**
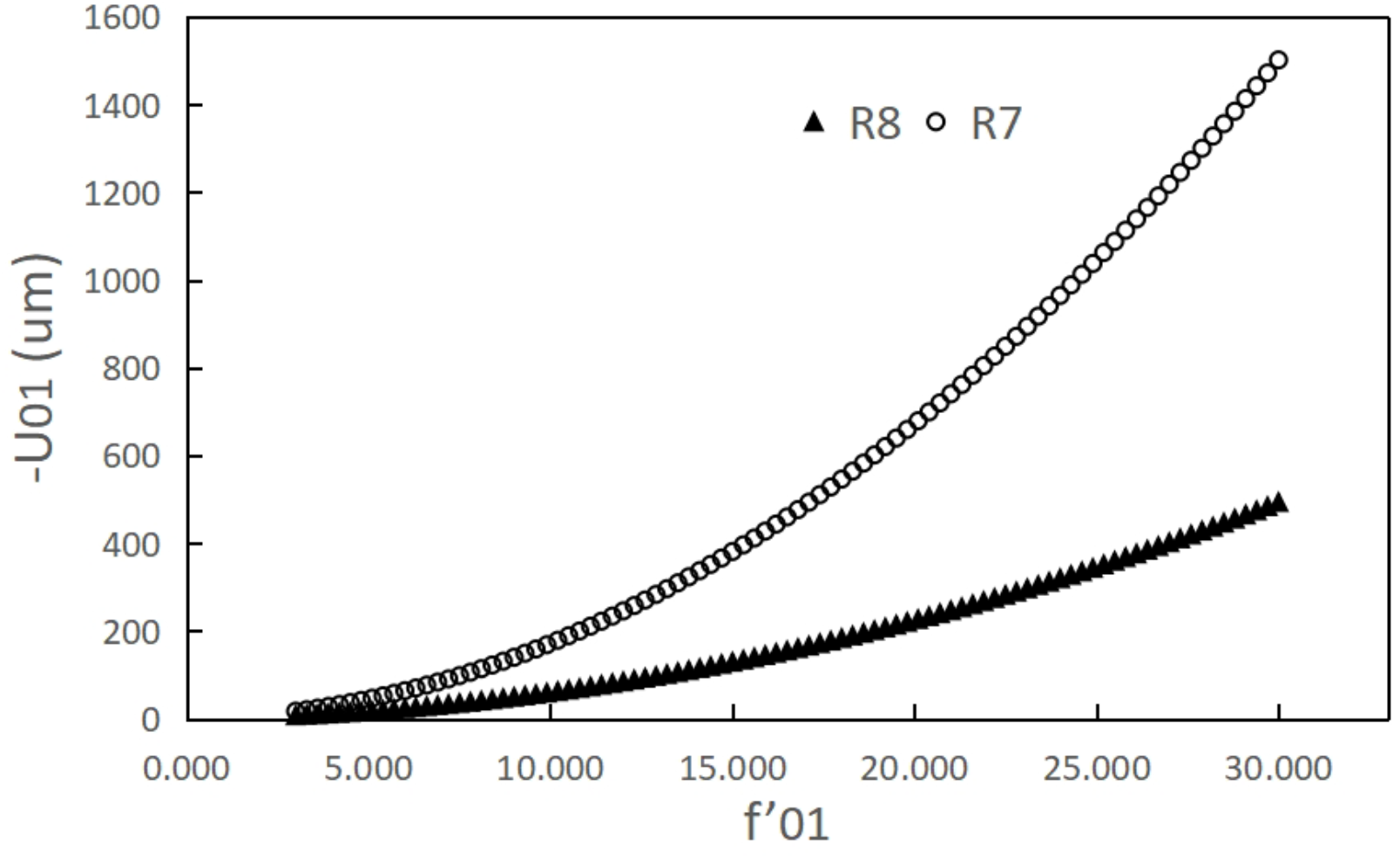
Depth perception simulation in *Drosophila* at Df = 0. With variable objective focal length f’0i = 3–30μm, object distances detectable by rhabdoms are: - R8: 6.96–493.0 μm - R7: 17.0–1501.3 μm Each microlens operates within distinct depth perception ranges when Df = 0.

**Figure 5.**
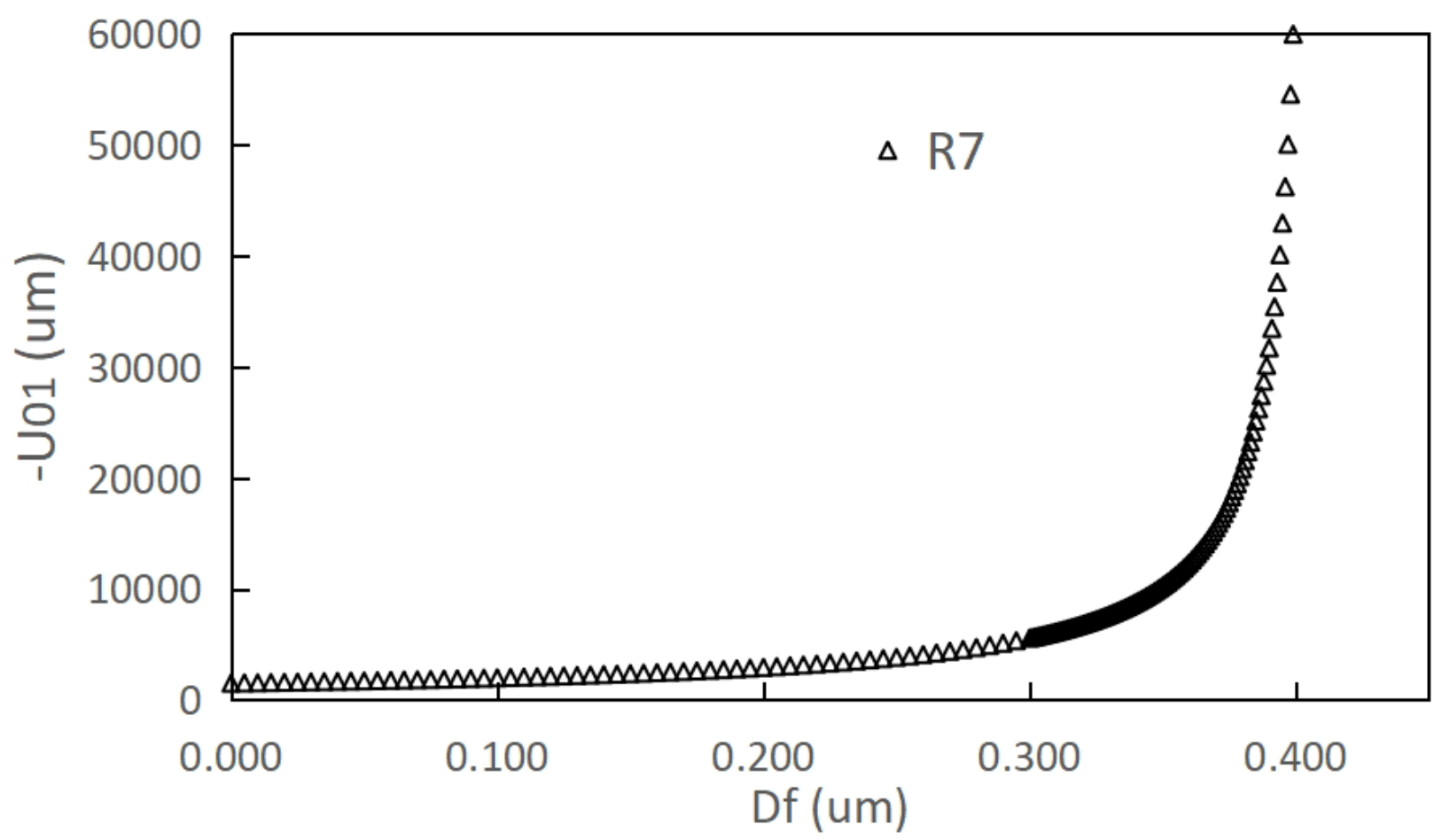
Depth perception expansion simulation in *Drosophila* via Df modulation. As (Df) increases (0–0.399 μm) with a fixed objective focal length of 30 μm, the detectable object range extends from 1500 μm (1.5 mm) up to ∼60 mm. Beyond Df > 0.409 μm, focal distance becomes infinite.

#### 2.4.3 Firefly Simulation

Using data from [7,8], extrapolated to larger-scale optics of fireflies (*Lampyris*) .

‐ Objective focal range: 10–200μm
‐ Df range: 0–0.181μm
‐ R7 detection range (Df = 0): =0.3–15 cm
‐ With Df expansion: up to 2.7 meters

#### Outcome

fireflies (*Lampyris*) , using single ommatidia, may resolve depth information across *orders of magnitude*, leveraging combined tuning of lens curvature and plane spacing.

**Figure 6.**
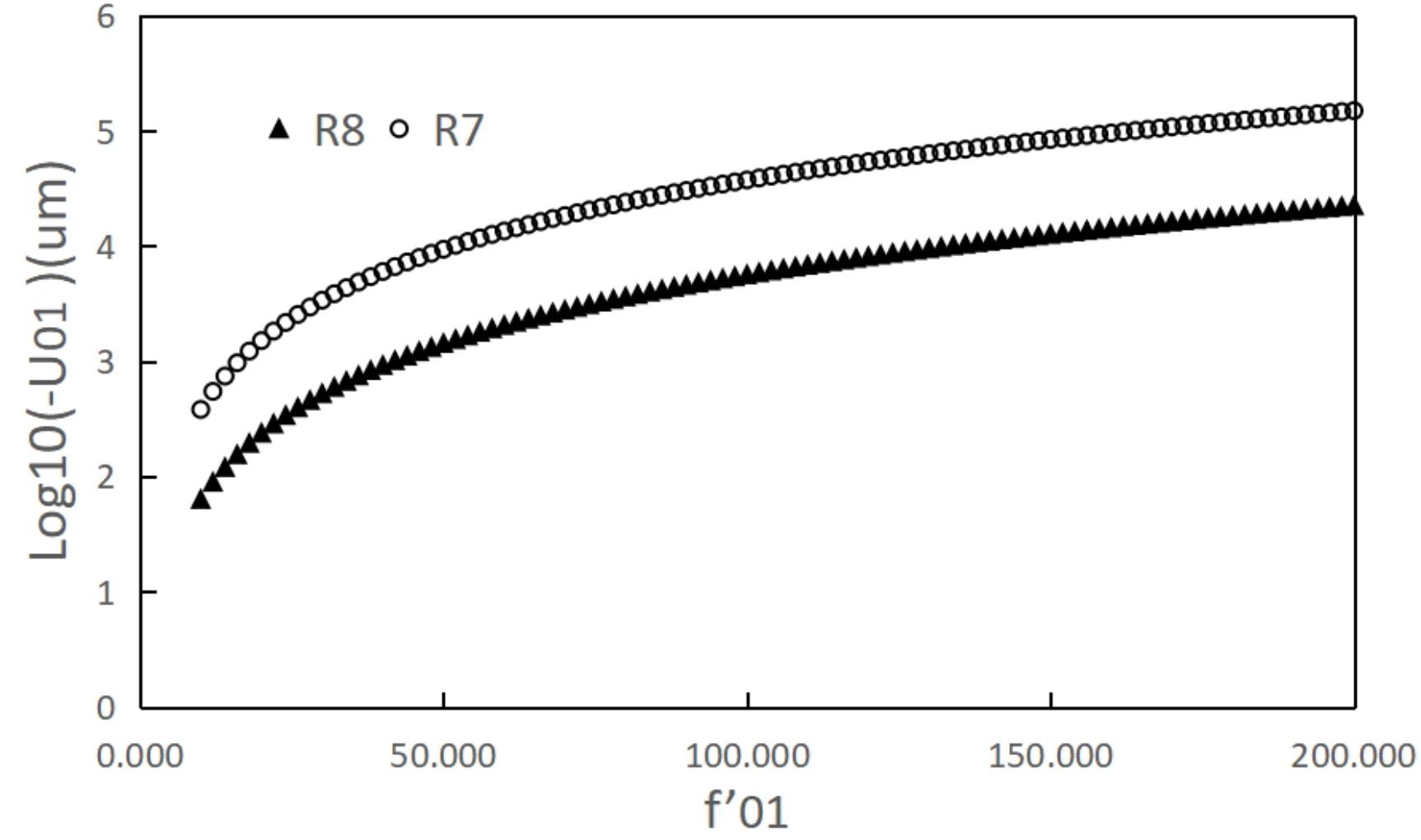
Depth perception simulation in fireflies (*Lampyris)* at Df = 0 (logarithmic scaling of distance) With focal length variation f’0i = 10–200μm, microlens depth ranges are significantly extended: - R8: 63.3 – 22,448 μm (∼2.2 cm) - R7: 381.8 – 149,863 μm (∼15 cm) This demonstrates the macro-scale depth sensing range of a single ommatidium.

**Figure 7.**
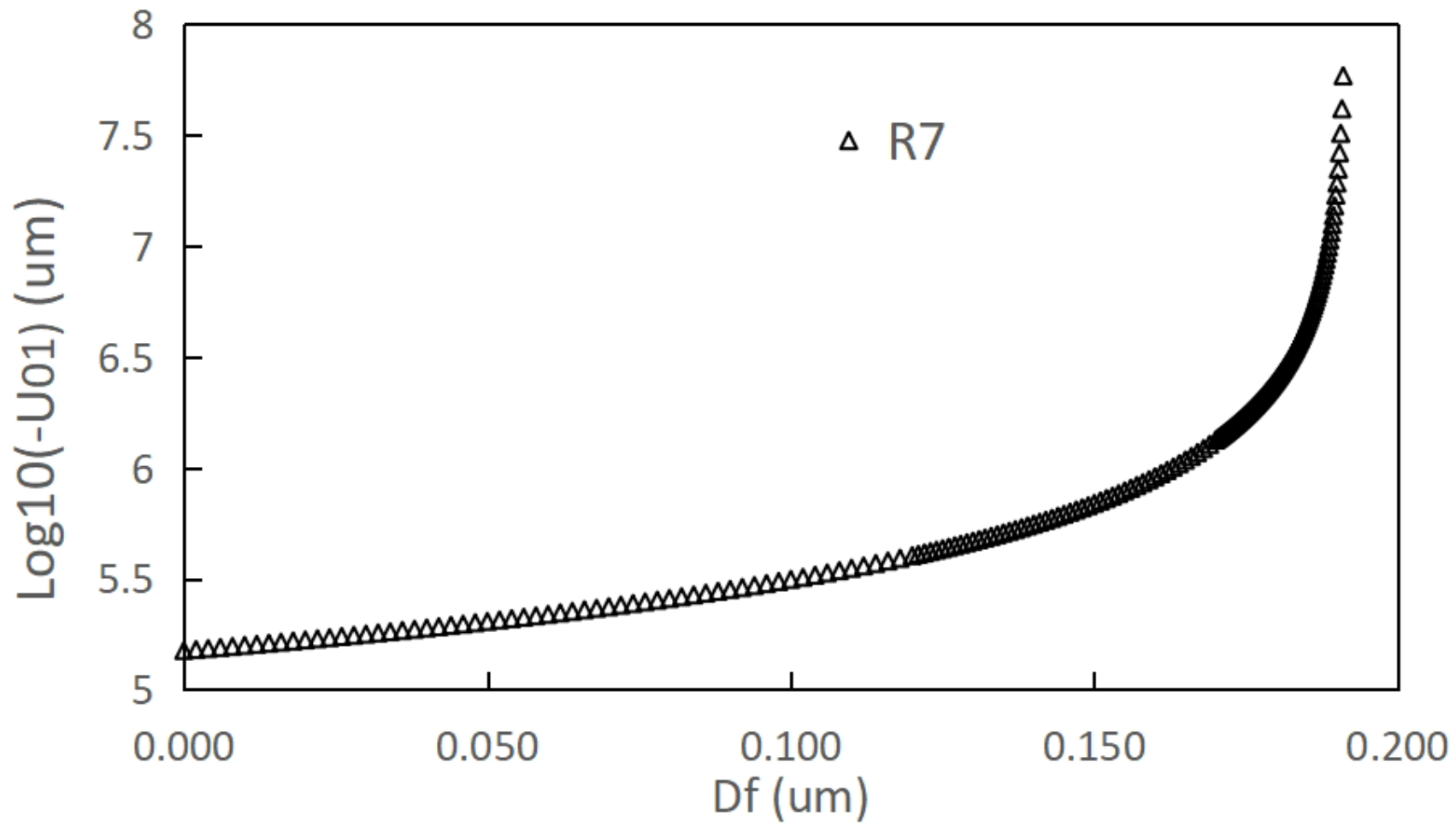
Firefly object distance sensing under increasing Df (focal separation), logarithmic scale. Keeping objective focal length fixed at 200 μm, increasing Df from 0–0.181 μm extends the object distance perception range from ∼15 cm to 2.7 meters. Df > 0.181 μm approximates focus at infinity.

## 3. Discussion

### 3.1 Feasibility of the 3D Optical Imaging Model in a Single Ommatidium and Reliability of the Simulation Results

The theoretical foundation of this study hinges upon a central hypothesis: **that the crystalline cones and rhabdomeric nuclei within a single compound eye ommatidium function as microlenses**. The optical viability of this idea depends on several biological and physical premises:

- **Density and refractive index:** Cell nuclei generally possess higher mass density than surrounding cytoplasm, implying a higher refractive index.
- **Size scale:** Given that nuclear diameters are several times larger than the wavelength of visible light, **geometrical (as opposed to wave) optics remain applicable**.
- **Simplifying assumptions:** In the present model, focal lengths and geometric arrangements of the microlenses have been idealized or inferred based on the literature. These assumptions have been made consistently across insect models for comparability and simulation accuracy.

While the model introduces a speculative but biologically plausible optical role for rhabdomeric nuclei, **experimental confirmation is essential to validate these proposed lensing effects and optical pathways**. Advanced microscopy (e.g., wavefront sensing, 3D tomography, live-imaging in genetically modified insects) could test and refine these predictions.

The simulation results based on estimated anatomical parameters and derived imaging equations suggest logical consistency in stereopsis mechanisms via distance mapping. Nevertheless, confirmation in vivo or in vitro will be key to establish the **biological realism of active microlensing at the sub-ommatidial level**.

### 3.2 Comparison to Traditional Apposition and Superposition Models of Compound Eye Imaging

Traditionally, compound eyes are classified into two primary imaging modalities [13]:

**1) Apposition optics (mosaic eye):**

- Each ommatidium perceives light from a specific directional cone, forming a single image point.
- Composite vision is constructed by combining signals across ommatidia.
- Spatial resolution is limited by inter-ommatidial angle and ommatidial number.

**2) Superposition optics:**

- Light from a single object point enters multiple ommatidia and converges via overlapping visual axes onto a central ommatidium.
- Typically seen in nocturnal species, allows greater photon collection.

In both models, each ommatidium acts as **a single-pixel detector**, registering just o**ne directional signal**.

#### Novelty of This Study’s Model

In contrast, **our proposed model posits that a single ommatidium can form multiple distinct image points** (up to four), and dynamically analyze depth information by engaging a **three-tiered imaging chain**:

1. **Primary image formation** — at first focal plane via objective (cornea + crystalline cone)
2. **Relay conjugate imaging** — via nuclear microlenses in upper rhabdom layer
3. **Final real-image focusing** — onto distinct rhabdomeric bundles (depth-specific photoreceptors)

This model effectively embeds a **stereoscopic optical device** within *each ommatidium*, permitting multi-angle, multi-depth analysis—exceeding the information density of conventional systems by a factor of 4.

### 3.3 Integration with Existing Theories of Insect Vision and Multi-Ommatidial Superposition

As illustrated in **Figure 8**, ommatidial overlaps known from classic superposition eyes may be augmented under our model—i.e., relay lens systems may function even across ommatidia:

- A central ommatidium (OMM(0)) may receive image contributions from neighboring units (OMM(−1), OMM(+1)).
- This implies a distributed but coherent three-stage relay across ommatidia:

**Figure 8.**
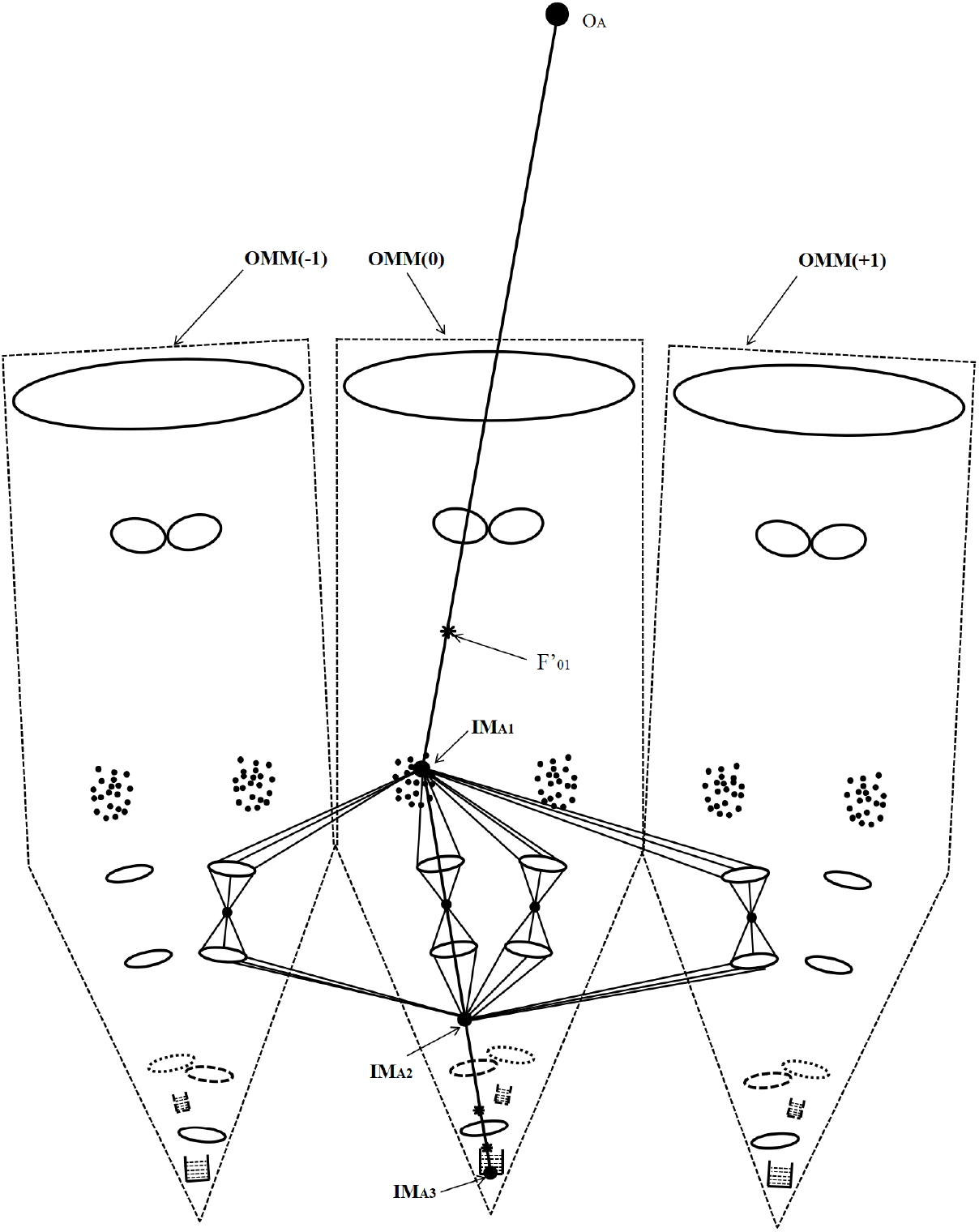
Schematic of multi-ommatidial overlap forming superimposed images. Caption: This diagram illustrates how multiple neighboring ommatidia (e.g., OMM(−1), OMM(0), OMM(+1)) may simultaneously capture light from a single spatial point and relay it toward the same inner rhabdom, forming a central overlapped image. Unlike traditional superposition eyes, this model utilizes an internal 3-stage imaging sequence: Primary image formation (first image, IMA1) Relay via conjugate mirror (IMA2) Final real image onto a rhabdom (IMA3)This suggests a higher efficiency use of input photons for depth perception.

▪ **Stage 1**: Directional light enters through discrete corneal sub-apertures.
▪ **Stage 2**: Multiple rhabdom nuclei act in parallel to conjugate first images— possibly even using inter-ommatidial nuclear arrays.
▪ **Stage 3**: Select microlens-rhabdom bundles, sensitive to depth, convert real images to electrical signals.

Such a **distributed compound eye-telescope array** theory offers a synthesis of apposition and superposition principles—and could enhance both resolution and depth estimation within massive, semi-autonomous ommatidial units.

### 3.4 Implications for Neurosensory Integration and Bio-Inspired Optical Systems

The results of this study offer several forward-looking applications and implications:

- **Biological significance**: This depth-sensing capability could explain **precise navigational and spatial behavior in insects** such as flight targeting, distance-based mating pursuit, and environmental mapping.
- **Neurological interpretation**: Signal spikes from rhabdoms of known depth positions (M7, M8, etc.) would serve as unambiguous cues for neural depth computation—effectively inferring 3D spatial coordinates from single-eye input.
- **Biomimetic design**: The “multi-angle real-image micro-telescope” model might inspire:
  ▪ 3D imaging sensors with **microscale variable focal depths**
  ▪ **Monocular auto-ranging** machine vision modules for drones or surgical tools
  ▪ High-resolution sensors with **nuclear-scale pixel units**

Combined optical-nuclear integration in this model represents a form of **subcellular optoelectronics**, where naturally self-assembled structures achieve functional imaging roles via emergent geometry and refractive contrast.

## 4. Conclusion

Based on the hypothetical microlens functionality of rhabdomeric nuclei within insect ommatidia, this study systematically proposes a new geometrical optical model whereby a **single ommatidium is capable of three-dimensional imaging and monocular stereoscopic perception**.

In contrast to conventional interpretations of compound eyes as arrays of flat, angular samplers of light (one image point per ommatidium), our model reconfigures each ommatidium into a **multi-angle, depth-sensing micro imaging system**. This is achieved by integrating:

- The **cornea and crystalline cones** as compound objective lenses for angular focusing;
- **Rhabdomeric nuclei** as coherent, directional **conjugate relay lenses**;
- Tiered **rhabdom bundles** as focal-depth-specific detectors.

With this three-stage system—**objective** → **relay** → **microlens**—we suggest that **insect brains could calculate object depth** by actively modulating lens parameters (either focal length or inter-focal plane distance) and decoding phototransduction events corresponding to image plane crossings.

Numerical simulations for three insect species (*Viggiani* wasps, *Drosophila*, and fireflies (*Lampyris*)) demonstrate that, using only one ommatidium and variations in focal parameters, the system can successfully detect **object distances ranging from micrometers to several meters**. These capabilities far exceed the constraints imposed by the traditional “one ommatidium, one pixel” paradigm.

This new model not only provides a theoretical framework for understanding spatial imaging and depth sensitivity in insects but also offers conceptual guidance for developing **biologically inspired imaging systems**, such as:

- **Microscale monocular 3D cameras**,
- **Multi-focal plane sensors** with nuclear-level pixel specificity,
- And **adaptive optics enhanced via structural approximations from biology**.

We hope this theoretical construct serves as a foundation for future **experimental validation and applied optical engineering**.

